# Improving data availability for brain image biobanking in healthy subjects: practice-based suggestions from an international multidisciplinary working group

**DOI:** 10.1101/110460

**Authors:** BRAINS (Brain Imaging in Normal Subjects) Expert Working Group, Susan D Shenkin, Cyril Pernet, Thomas E Nichols, Jean-Baptiste Poline, Paul M. Matthews, Aad van der Lugt, Clare Mackay, Linda Lanyon, Bernard Mazoyer, James P Boardman, Paul M Thompson, Nick Fox, Daniel S Marcus, Aziz Sheikh, Simon R Cox, Devasuda Anblagan, Dominic E Job, David Alexander Dickie, David Rodriguez, Joanna M Wardlaw

## Abstract

Brain imaging is now ubiquitous in clinical practice and research. The case for bringing together large amounts of image data from well-characterised healthy subjects and those with a range of common brain diseases across the life course is now compelling. This report follows a meeting of international experts from multiple disciplines, all interested in brain image biobanking. The meeting included neuroimaging experts (clinical and non-clinical), computer scientists, epidemiologists, clinicians, ethicists, and lawyers involved in creating brain image banks. The meeting followed a structured format to discuss current and emerging brain image banks; applications such as atlases; conceptual and statistical problems (e.g. defining ‘normality’); legal, ethical and technological issues (e.g. consents, potential for data linkage, data security, harmonisation, data storage and enabling of research data sharing). We summarise the lessons learned from the experiences of a wide range of individual image banks, and provide practical recommendations to enhance creation, use and reuse of neuroimaging data. Our aim is to maximise the benefit of the image data, provided voluntarily by research participants and funded by many organisations, for human health. Our ultimate vision is of a federated network of brain image biobanks accessible for large studies of brain structure and function.

## 1. Introduction

Neuroimaging has become embedded in substantial research endeavours to understand normal brain function and effects of disease (e.g. Thompson et al., 2003; Fox and Schott, 2004; Lemaitre et al., 2005; Marcus et al., 2009; Wardlaw et al., 2011; Weiner et al., 2016). Until recently, many neuroimaging studies were in single centres and, inevitably, of modest size (Dickie et al., 2012). Many much larger population scanning initiatives are now ongoing (Jack Jr et al., 2008), and many multicentre clinical trials routinely include imaging as part of inclusion criteria and as outcome measures (Cash et al., 2014), providing the potential for large multicentre collections capturing the range of brain structure in the population. The importance of maximising the value captured in this large amount of imaging data – to detect how differences in brain structure and function relate to behavioural or clinical outcomes – is now widely recognised (Toga, 2002; Barkhof, 2012; Poline, 2012). The value of data for answering new questions can grow with sample size, e.g. for replication, increasing population representativeness, and increasing study power. To address this issue, a growing number of electronic databanks including brain imaging are available, either from dedicated cohorts (e.g. Alzheimer’s Disease Neuroimaging Initiative, UK Biobank, IMAGEN), or collections of studies (e.g. Brain Imaging in Normal Subjects, Dementia Platform UK, Open Access Series of Imaging Studies): see Table 1.

**Table 1.** Databases presented at brain image bank meeting with relevant references, website, and data access policy

| Data Base | Website | Data access policy |
| --- | --- | --- |
| ADNI (Weiner, 2015)<br>ENIGMA (Thompson et al., 2015) | http://adni.loni.usc.edu/<br>http://enigma.ini.usc.edu/ | ADNI: registration and application for approval by steering committee for data access<br>ENIGMA: data is shared between members on an ad hoc basis |
| BRAINS (Job et al., 2016) | http://www.brainsimagebank.ac.uk/ | Open access search of available data; registration and |
|  |  | application for approval by steering committee for data access |
| Dementia Platform UK (DPUK) | http://www.dementiasplatform.uk/ | In development: online registration required |
| Edinburgh Birth Cohort | www.ebc.ed.ac.uk | Online search of available data ; registration and application for approval by steering committee for data access via http://www.brainsimagebank.ac.uk/ |
| European Population Imaging Infrastructure Rotterdam study (Ikram et al., 2015 ; Hofman et al., 2015)<br>Generation R (Jaddoe et al., 2012) | http://populationimaging.eu/<br>http://www.erasmus-epidemiology.nl/research/ergo.htm<br>http://www.generationr.nl/ | Email contact for information on available datasets |
| EVA, 3-CITIES, DBGIN, BIL&GIN, i-SHARE (Alperovitch et al., 2002; Lemaitre et al., 2005; Mazoyer et al 2016). | http://www.gin.cnrs.fr/BILandGIN<br>http://www.three-city-study.com/the-three-city-study.php | Registration and application for approval by steering committee for data access |
| IMAGEN | http://www.imagen-europe.com/en/consortium.php | Registration and application for approval by steering committee for data access |
| Montreal Consortium for Brain Imaging Research (Evans et al., 2012). | https://www.mcgill.ca/globalhealth/international-consortium-brain-mapping | Registration and application for approval by steering committee for data access |
| OASIS (Marcus et al., 2007b, 2009)<br>Human Connectome Project (Van Essen et al., 2013) | http://www.oasis-brains.org/<br>http://www.humanconnectome.org/ | OASIS: open access, no registration required<br>HCP: open access with restricted access for sensitive data (registration and application for approval by steering committee) |
| Rhineland study | https://www.dzne.de/en/research/research-areas/population-health-sciences/rhineland-study.html | Email contact for information on available datasets |
| UK Biobank (Matthews & Sudlow, 2015) | http://www.ukbiobank.ac.uk/ | Open access search of available data; registration and application for approval by steering committee for data access |

### Brain images from ‘healthy’ subjects are important

The wide variation in brain structure and function both within and between individuals at different ages has long been recognised (Wardlaw et al., 2011; Dickie et al., 2013). Methodologies that use appropriately representative populations are needed to provide normative populations, particularly for healthy subjects (i.e. those without neurological diseases such as stroke or dementia). They can provide informative reports for users (e.g. ‘brain on 5^th^ percentile for volume at age 70’ for a specified population) and simultaneously embrace the spectrum of individual variation (Dickie et al., 2015a, 2015b). Brain imaging is increasingly used in the diagnosis of neurological diseases, and mental health disorders (Fox and Schott, 2004). Data from existing cohort or population studies (e.g. Marcus et al., 2009), can help define boundaries between health and disease, to aid diagnosis and trial inclusion, to provide effect size estimates for planning trials, and, where relevant, controls for case-control studies (e.g. Dickie et al., 2015a; ADNI: Potvin et al., 2016).

### Current status of brain imaging banks

Large repositories of brain imaging data from well-4=lrwiop[/[/]74characterised subjects in accessible databanks are required to achieve this, while ensuring that data protection concerns are also addressed. These comprise data initiatives that are planned around harmonised protocols, such as ADNI (Alzheimer’s Disease Neuroimaging Initiative) (Weiner et al., 2015), UK Biobank (Matthews & Sudlow, 2016), Human Connectome Project (van Essen, 2013), OASIS (Open Access Series of Imaging Studies) (Marcus et al., 2007a, 2007b, 2009), and those that represent data aggregation without initial harmonisation e.g. ENIGMA

(Enhancing Neuro Imaging Genetics through Meta-Analysis-Thompson et al., 2014, 2015). The value of brain images is hugely enhanced by the information on the characteristics of individual subjects and the study in which they participated, but at present studies vary widely in what data they present on the study, subject or image data, and how these data are presented (Dickie et al., 2012).

Only a small proportion of the images performed for research are included in biobanks, and in existing structural brain image biobanks, normal subjects over 60 years of age are relatively under-represented, with limited cognitive and medical metadata to support their classification as “normal” (Dickie et al., 2012), and available with a limited range of neuroimaging sequences. For example, fluid attenuated inversion recovery (FLAIR) and T2* volumes are often not available, although they are essential for sensitively identifying and quantifying white matter hyper-intensities (WMH) and microbleeds respectively, neuropathologies present in normal ageing but associated with vascular cognitive impairment (Wardlaw et al., 2013; Ritchie et al., 2016). Newer initiatives like BRAINS (Job et al., 2016) provide a range of sequences (e.g., T1, T2, T2*, and FLAIR) for most subjects plus cognitive and medical information. Future data sharing will be facilitated by influencing how new data are collected in terms of core imaging sequences and meta-data variables.

### Standards for sharing

The INCF (International Neuroinformatics Coordinating Facility) Standards for Data Sharing Neuroimaging Task Force the Brain Imaging Data Structure (http://bids.neuroimaging.io/) to advance standard organisation and descriptions of data files, and the Neuroimaging Data Model (http://nidm.nidash.org/) for data provenance tracking, but ongoing work is needed around developing community consensus and adoption of standards (Bjaalie and Grillner, 2007). Issues such as privacy, de-identification, quality control, provenance, avoiding including the same subjects in multiple databases, ethics (historical and future), consent, essential components of ‘good guardianship’, costs, sustainability, software version control, definitions of ‘normality’, and international variations in ethical and legal frameworks, also need further consideration (Rodríguez González et al., 2010). The European Society of Radiology (ESR) published a position paper on Imaging Biobanks (European Society of Radiology, 2015) defining imaging biobanks, outlining their purpose, and advocating the creation of a network/federation of such repositories with existing biobanks.

Many funders advocate or mandate that data generated by studies they fund are made public and the International Committee of Medical Journal Editors (ICMJE) has proposed that deidentified patient information is shared before research manuscripts of randomised controlled trials will be considered for publication (Taichman et al., 2016). While this data sharing may be relatively straightforward for tabular demographic data (i.e. the types of alphanumeric data that can be held in traditional databases), the situation is much more complex for brain image data (Toga, 2002, Marcus et al., 2007a). Factors like the size of imaging files and the possibility of identifying subjects from images impose non-trivial technological challenges. While initiatives such as NeuroVault (http://www.neurovault.org -Gorgolewski et al. 2015) avoid the problem by publicly sharing statistical maps for data aggregation it does not include whole datasets. By contrast, a repository like OpenfMRI (http://www.openfmri.org) includes raw-data, with some subject-level variables, which allows newer analyses to be performed. Even when there is a desire to share imaging data, there are a number of technical, legal and practical problems to be overcome: (Poline 2012; Poldrack and Gorgolewski, 2014, Pernet & Poline, 2015).

## 2. Learning from existing databanks and population studies

Against this background, a group of experts, including specialists in image acquisition and analysis, clinical disciplines, epidemiology, legal, ethics, and data science, met to discuss and debate conceptual, legal, ethical and technical issues around creating brain image banks. We aimed to highlight the issues that need to be addressed, from the ethical to the practical, achieve some consensus, promote best practice and provide useful advice for ongoing and planned studies. The primary aim of the meeting was to encourage data sharing, construct pan-institutional brain image databank consensus, and facilitate linking between databanks. Here, we describe lessons learned from existing image databanks, provide advice on technical, epidemiological and legal challenges and identify areas where agreement was not reached that should be addressed as the field evolves.

Representatives of major groups involved in neuroimaging databases were present at the meeting (Table 1). We recognise that there are other imaging databases, many summarised in recent publications (e.g. Sharing the Wealth: Brain Imaging Repositories in 2015; Neuroimage 124 Part B), but here we provide the lessons learned by this international group with experience in building databases and sharing data under various schemas, for healthy participants and people with neurological disorders, from prenatal to old-age. We noted that the information collected in each databank was very different depending on the perspective and expertise of the individual, e.g. computer scientists or data analysts versus clinicians or epidemiologists.

Among the various problems that plague databases, the group identified four, which raise new questions not previously well addressed in the epidemiological community for neuroimaging: (1) data collection; (2) addressing data heterogeneity; (3) database infrastructure; and (4) database management. These aspects marks major divisions that goes into building a databank (Figure 1). Here, we discuss each of these aspects, presenting lessons learned and recommendations.

**Figure 1.**
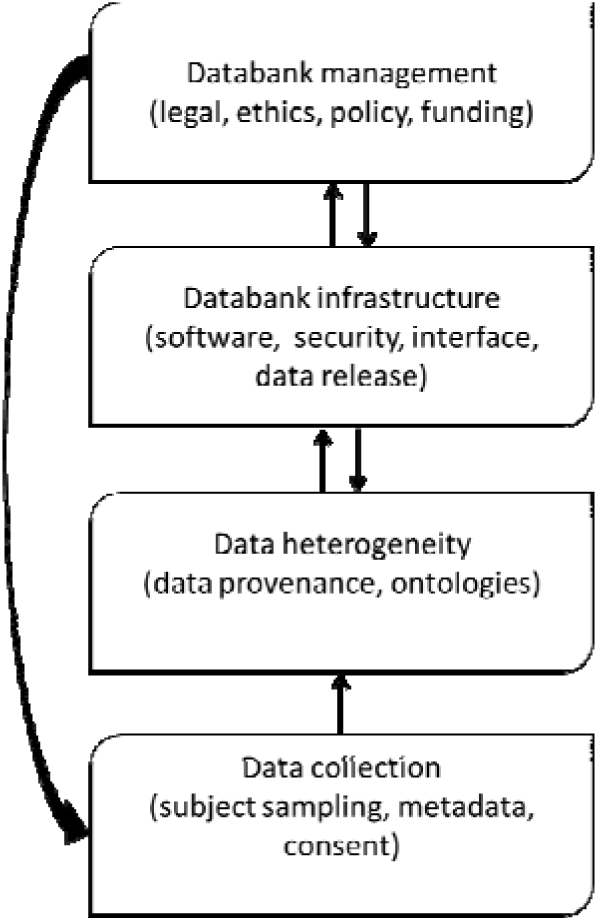
Major issues identified in building a brain image biobank (arrows indicate which aspects constrain each other)

## 3. Data collection

There is a great willingness from many people across the life course to volunteer for brain imaging studies: even when the participants are in their nineties and the study includes prolonged imaging (Deary et al., 2012). However, such willing individuals – irrespective of age – tend to be fitter, better educated and less socially deprived than the general population (e.g. Deary et al., 2012; Stafford et al., 2013). Extra effort is therefore needed to encourage more representative population sampling, or the consideration of statistical approaches to account for known bias (Ganguli et al., 2015).

### 3. A. Subject sampling

Several aspects of study design, fundamental to good epidemiology, are currently not prominent among brain image banks. The use of guidelines on study reporting should improve quality (http://equator-ntework.org). The method of subject selection (random, sequential, particular characteristics, etc.), population from which they were selected (e.g. hospital clinic attendees, primary care attendees, general population) and method (e.g. direct mailing, adverts, any compensation or payment) should be described clearly. In general, research participation varies with social class, education, health status, by ethnic and other minority groups (Deary et al., 2012; Stafford et al., 2013), hence documenting the original study aims when data are collated for secondary use is important to describe the population. This can be achieved by citing a paper, ‘read me’ descriptors, or website that describes the original study, linked from the database.

A subject should only contribute once to a particular analysis. However, several open datasets include the same individuals (Dickie et al., 2012) potentially distorting any results if several such databanks are used in one analysis. Methods are needed to identify if a subject is included in multiple studies. Identification of the uniqueness of a subject’s inclusion in a database is a significant problem, with few effective solutions at present. ‘Pseudo– anonymised’ identifiers can identify individuals in longitudinal studies. Probabilistic matching of clinical or imaging data could identify repeat subjects if enough data were available – however this approach may not be reliable, and also implies that even de-identified data about an individual cannot be considered truly anonymous, as data may be susceptible to data linkage attacks (Fung et al., 2010). Simply asking participants about inclusion in other studies is unlikely to be reliable. Preferably, subjects would be assigned a unique study identifier linked to a unique study registration, but this is difficult to implement in practice, as there is no central database of studies. However, tools such as The Global Unique Identifier (GUID) may perform this function (https://data-archive.nimh.nih.gov/rdocdb/s/guid/nda-guid.html). GUID is a universal subject ID which allows participants to be matched across labs and research data repositories, as well as allowing researchers to share data specific to a study participant without disclosing identifiable information. In genetic studies, it is sometimes possible to use a “checksum” method based on a person’s genomic data, to test whether any individuals took part in two studies being compared, as this may be important to avoid reporting spurious associations (Franke et al., 2016).

Methods for long term follow up of healthy volunteers and adding linked data including health outcomes many years later, is potentially very valuable and should also be considered wherever possible (e.g. Deary et al. 2011; Matthews & Sudlow, 2015).

### 3. B. Metadata collection

Some studies include extensive socio-demographic data, others have detailed clinical characteristics and vascular risk factors, others focus on cognitive testing, and several have extensive blood or urine biomarkers including genetics. A key aspect to consider is to define the minimum subject metadata to ensure maximum use of the data world-wide.

**Age, sex** (self-report)**, handedness** (self-report and/or Edinburgh inventory test) and **education** (total years of education and highest qualification) are important variables that should always be reported, because they are associated with brain structure and function: age (Dickie et al, 2012); sex and handedness (Good et al, 2001); and education (Cox et al, 2016). The detail in which variables are collected, and what other meta-information is required depends on the setting. In a clinical setting, ‘normal’ or healthy is often defined as above or below a cut-off, the definition may be somewhat arbitrary (e.g. blood pressure of 140/90; cognitive test score above a threshold; blood test value). In a research setting, the concept of ‘normal’ may refer to healthy controls, who do not have the disease of interest (but may have other conditions) and meet several other criteria defined within the study. People recruited to a study because they are ‘healthy’ or ‘normal’ may have occult or undeclared disease, have borderline values (e.g. blood pressure of 138/88) or develop disease in the future. It is essential that those participating in a study are well-described if their data are to be used for other purposes. A balance needs to be struck between overburdening subjects, versus inadequate information. The method of collection of data (e.g. self-report of sex vs. chromosomal identification) will depend on the purpose and location of the original study, and description of the population is encouraged (e.g. educational system to allow international comparison).

Further discussions are required to agree a minimum normative subject dataset, but this is likely to include some measure of **sociodemographic variables, clinical features (main comorbidities, medications), cognitive ability (at least a general cognitive test)**.

Availability of **biomarkers** (such as DNA) may also be useful. The potential to **link** to data collected for other reasons, such as participation in nationwide Biobanks or data collected during clinical care, may reduce the burden on participants if appropriate consent is provided and relevant reliable data are available.

### 3. C. Consent

Individuals must be informed and provide informed consent where possible. One issue is that it may be impossible to re-contact previous participants who have already provided consent for further use of their data, where imaging databanks were not specifically mentioned. In such case, retrospective approval might be given by an ethics committee. More importantly for new studies, consent must include information on data reuse and sharing. The Open Brain Consent initiative (https://open-brain-consent.readthedocs.io/en/latest/) provides sample consent forms to allow deposition of anonymised data to public data archives, and a collection of tools to facilitate anonymization of neuroimaging data to enable sharing. This initiative aims to facilitate neuroimaging data sharing. It is based on the legal and ethical framework of the USA, but is adaptable to other countries.

Individual projects should also decide whether there is a need to link subjects back to their anonymised data, and consent therefore must include information on being re-contacted. Individuals and researchers should be informed of who ‘owns’ the data (often the host institution: principal investigators should be aware of any restrictions if they move between institutions), if there are copyrights on images, and consent could be indicated in a metatag. It is also important to consider what should be done if data are collected from people without capacity to consent (e.g., children: if consent is provided by a parent on their behalf, can the child rescind this on reaching adulthood, and when should a young person be asked to give consent to ongoing participation in a longitudinal study that his / her parent consented to during the child’s infancy).

### 3. D. Practical consideration in collecting MRI scans across the lifespan

#### Perinatal

Data from healthy ‘normal’ fetuses throughout gestation and neonates born at term are currently sparse. Single centres have generally developed repositories to study specific cohorts (e.g. congenital heart disease or preterm neonates), using bespoke processing pipelines or pipelines developed using adult data but optimised for neonates (Boardman et al., 2006; Miller et al., 2007; Ball et al., 2010; Serag et al., 2016). There are some fetal and neonatal structural MRI atlases (for example: Gousias et al., 2012; Shi et al., 2011; Blesa et al., 2016; Makropoulos et al., 2016; Kabdebon et al., 2014; Oishi et al., 2011), but to our knowledge there are no perinatal image banks hosting normal data acquired from multiple studies and sites. A perinatal sub-section of the BRAINS database is under development, and the developing Human Connectome Project (http://wp.doc.ic.ac.uk/dhcp/) aims to make data available from 1500 fetuses and newborns between 20-44 weeks’ post-menstrual age.

Experience from perinatal brain repositories is that newborn neuroimaging research is acceptable to parents and carers, especially those with children at risk of long term impairment. Newborns, including those who require intensive care, can be looked after safely in the MRI environment (Merchant et al., 2009). Research quality data can be acquired from infants without the use of sedation by using feed-and-wrap techniques, and by allowing flexibility within in the scan schedule to allow for coaxing to sleep and managing wakefulness. Perinatal image data are readily contextualised by maternal, pregnancy and birth information, and it can be analysed with information from infant biosamples (Boardman et al., 2014; Sparrow et al., 2016) and standardised childhood neurodevelopmental / cognitive outcome tests (Woodward et al., 2006): achieving prospective consensus over minimum datasets in these domains will maximise opportunities of perinatal data-sharing in future initiatives.

#### Older age

Normal ageing is associated with increasing variability in brain structure, such as brain tissue loss (atrophy) and the accumulation of white matter hyper-intensities, which may or may not have functional impact on cognition, mood, gait etc. (Dickie et al., 2015a). Many brain image biobanks were created to study pathological ageing, e.g. dementia, and only small numbers of ‘healthy’ older people were previously included in accessible databanks (Dickie et al., 2012), making it difficult to define ‘normal ageing’ or pathological ageing.

The availability of data from UK Biobank (http://www.ukbiobank.ac.uk/) and other ongoing population imaging initiatives in North America and mainland Europe will change this. One conclusion from the workshop was that, for databanks to accurately represent the spectrum of health, a large number of participants, sampled in equal numbers across appropriate age bands are required, accompanied by detailed descriptions of the demographic and clinical characteristics of individuals.

## 4. Addressing Data heterogeneity

Where more than one study is included in a brain image bank, like 3-CITIES (Alperovitch et al., 2002) or BRAINS (Job et al., 2016), there is usually substantial heterogeneity of the acquired demographic/clinical and imaging data. This can be addressed either by describing each variable (3-CITIES), or by harmonising metadata (BRAINS). Having many variables makes the database large and difficult to search, while transforming variables to agreed standards, which is simpler for the end user, is resource intensive and requires additional data provenance documentation. Alternatively, in approaches such as ENIGMA (Thompson et al., 2015) and UK Biobank (Matthews and Sudlow, 2015), both raw and derived data from each of the participating centres can be used for large scale analyses: either in meta-analyses, which can circumvent issues of data sharing and transfer between countries, but may restrict the analyses that can be performed; or in ‘mega’-analyses using original data, which may be difficult to access due to data access controls, and/or too different to combine. Choosing the right framework clearly depends on the question(s) the database aims to address, and requires collaboration between local data providers and the databank.

### 4. A. Minimum provenance of study data

Brain imaging uses indirect measures to make inferences about brain structures. The meaningfulness of these measures will vary with the amount and heterogeneity of the data. The version of scanner and software used for data collection and analysis should be clearly documented. Decisions need to be made about the inclusion of raw or processed data in the database, recording of processing steps, and whether outputs of imaging data will be raw, pre-processed or fully processed. The amount of data storage required for imaging (and e.g. genetic) data may require the use of high performance networking, storage and computing, which can be upgraded without compromising the database, adequate bandwidth, and/or the use of cloud computing, taking account of privacy and security issues (Poldrack 2014).

Data provenance (study, aims, date performed, funders, principal investigator, recruitment method, publications) is important for appropriate citation and recognition of data sources, encouraging reproducibility, avoiding duplicates, and data versioning. This information can easily be documented on a per study basis without complex tools, though formal representation will maximise the potential for reuse. For example, the W3C-PROV specifications is a framework to interchange provenance information (http://www.w3.org/TR/prov-dm/), and its extension for neuroimaging, NIDM (http://nidm.nidash.org -Keator et al., 2013), provide a way to encode provenance in a machine-readable manner. The European Human Brain Project (https://www.humanbrainproject.eu/enGB) is also developing guidance on best practice for data mapping and sharing.

From a technical perspective, it is important that image related information are all recorded and shared. The Organisation for Human Brain Mapping recently provided a **consensus list of reporting items** (which can be a useful starting point for collecting and harmonising site-specific information on imaging data (see appendix D, especially table D2 - http://www.humanbrainmapping.org/COBIDAS; Nichols et al., 2016).

### 4. B. Role of Ontologies

To allow comparisons between banks, many groups are now working on methods to compare clinical and imaging variables, and using appropriate ontologies. General standards can be found in the NIH’s Common Data Elements (http://cde.nih.gov/), an attempt to collect terminologies across biomedical practice. Building on PROV-DM (http://www.w3.org/TR/prov-dm/) - a World Wide Web Consortium standard to describe provenance - a Neuroimaging Data Model (NIDM) has been developed by the Standards for Data Sharing Neuroimaging Task Force of the International Neuroinformatics Coordinating Facility (Keator et al., 2013) to provide a unified framework on image format, names and image meta-data. XCEDE (XML-extensible markup language-based Clinical and Experimental Data Exchange: Gadde, 2012) provides general standards for data management, but specific terms must also be used. Subjects’ metadata are much more difficult to describe and cognitive tasks can be particularly difficult to aggregate or compare meaningfully. A recent attempt to describe them has been made in the Cognitive Paradigm Ontology (Turner, 2012) or the Cognitive Atlas (Poldrack et al. 2011). It is important that the databank clearly describes which ontology was used, how decisions were made, and that all metadata variables are clearly defined.

## 5. Database infrastructure

Many of the studies that led to the creation of imaging databanks started over a decade ago, and reported issues relating to changing technology (Mazziotta et al., 2001). For example, technical staff need to consider the impact of hardware changes (e.g. upgrading or changing scanner software or hardware; changes in data storage solutions and formats) and software evolution, which can make keeping track of multiple analyses of the database challenging (Poldrack, 2014). Such changes in technology have, for instance, been shown to impact on local brain volume (Lorio et al., 2016) and atrophy measurements (Leung et al., 2015).

Changing requirements for data governance and data management also need to be considered: file names and structures may need to be updated to newer recommendations such as BIDS (Brain Imaging Data Structure) (Gorgolewski et al., 2015). Similarly, better understanding of disease aetiology, or changes in taxonomy, may affect how clinical characteristics are coded, and therefore what a database entry means, e.g. changing definitions of dementia subtypes. Studies may be cross-sectional or longitudinal. In studies of development or ageing, longitudinal data are particularly valuable (Mills and Tamnes, 2014), and systems need to be in place to ensure that future data acquisition can be matched to the correct subject, and that imaging parameters are similar enough to allow comparison.

### 5. A. Technical infrastructure

To maximize usage and usability, any planned databank should make use of a formal imaging database tools. We considered five software tools that create sharable, searchable databases and offer maximum flexibility: COINS (http://coins.mrn.org/) (Landis et al, 2016); LORIS (http://mcin-cnim.ca/neuroimagingtechnologies/loris/) (Das et al 2012); NiDB (https://github.com/gbook/nidb); Scitran (http://scitran.github.io/); and XNAT (http://www.xnat.org/) (Marcus et al, 2007a). The ability to link and search make those applications different from repositories; however each has strengths and limitations, some significant. Information reported below reflect user experience and discussion with software developers.

Each of these different software tools has different strengths, discussed at the meeting. Comparing first the aspects related to the nature and format of **imaging data,** all of the above software tools support the Digital Imaging and Communications in Medicine (DICOM - http://dicom.nema.org/) format and can interface directly with scanners. Some software tools support almost any other image formats (COINS, NiDB, XNAT), while others are restricted to specific ones (LORIS, Scitran). Only XNAT has an explicit tool to link the data from the database directly to clinical PACS (Picture Archiving and Communication System), i.e. allowing direct transfer between the database and the health care service. As imaging is just one facet of any research study, it is vital that all these tools also allow storing of **non-MRI data** such as demographic, clinical, behavioural, and genetic data. Other types of data such as electroencephalography can be stored but may not be viewable using imaging analysis/display interfaces, and may require additional software.

Another important aspect to consider relates to the **software**: (1) How are MRI data linked with other types of data? (2) How can the data be visualised?; and (3) How can the data be explored? Linkage (bringing together datasets) is an important aspect to consider regarding the size of the dataset and its utility. For instance, LORIS is built with two distinct databases (imaging vs non-imaging) that allows the user to interrogate, process and retrieve information separately or together. XNAT uses an XML-defined schema for searching data, and also uses and underlying object-relational database PostgreSQL. Like LORIS, this allows searching and accessing data separately. Considering **the end-user perspective** of the database one wishes to build, all five tools allow one to search for image data within and across projects and to visualise data, though with some restrictions depending on the software.

The final important aspects to consider are the overall **maturity, usability, maintainability, extensibility and support** of the software and **access control**. LORIS and XNAT, for instance, are well established and maintained, while newcomers (COINS, Scitran, NiDB) are more adapted to new data types and use more recent software technology. Mostly, the software can be installed with minimal programming expertise – but this differs between tools. Setting up multiple access control levels, linking or adding new functionalities, can however require much more expertise. For instance, XNAT, the most widely used platform, requires programming expertise to enable its extensible XML structure. In contrast, simpler tools with extensive search solutions exist (e.g. LORIS, NiDB), but modifying the search tool to suit dedicated needs also requires dedicated programming expertise. In terms of usability, it is worth considering data visualization, and whether the pipeline includes data quality control or basic data pre-processing. All the platforms considered include such options but various levels of programming expertise are required. It is important to also consider ‘futureproofing’ the technology, at a minimum ensuring accurate version control, and direct access to code. Another essential aspect to consider is **access control**: COINS, LORIS, Scitran, XNAT have extensive security levels to create, read, and edit data, while NIDB is limited to user/project management. Thus, at present, the consensus was that there is no ‘best’ software, the field is moving rapidly, and it is worth considering every aspect discussed here with expert advice to choose a software tool that best suits specific needs.

### 5. B. Security

A brain scan is unique, and could allow identification of the individual (e.g. by someone who already holds a copy of their brain image). There is a trade-off between removing all potential identifiers and retaining the scientific value of the data. Clear processes are required regarding sensitive, or potentially identifiable, data to ensure that all reasonable safeguards are put in place, e.g. the DICOM Confidential software; completely removing all textual personally identifiable information; generating new “anonymous” identification numbers; “defacing” brain MRI (Marcus et al., 2007b, Milchenko and Marcus, 2013; Rodriguez, 2010). A balance between accessibility and security is important, ensuring that potentially identifiable data (text or images) are protected from remote access in the database and if released for external use. It should be clear to the intended users what the data are, and what will be the safeguards to access. To test whether data can be hacked, a mock database can be developed, released and attempts made to infiltrate the security systems.

### 5. C. User interface

The user interface should suit the main proposed users. The requirements of clinicians, researchers, or industry are however likely to be very different (for instance searching per pathology vs. scan feature, looking for raw vs. processed data). It is worth considering the user interface and its design. In XNAT for example, the level of detail displayed to users can be made dependent on both their relationship with the databank (external user, contributor, etc.) and/or the intended use of the data of interest (Marcus et al., 2007a). The use of an Application Program Interface (API) allows for easier creation and customisation of user interfaces for different user groups, but can bring new security concerns.

### 5. D. Data release

In ongoing/longitudinal studies, there is a wide variation in when data are released: as collected, in batches, on completion, or after all analyses performed by initial research team. There is currently no consensus on how to release data, but it is important to make that decision clear and have mechanisms in place to identify releases (data version control). Primary researchers who developed the database should apply through the same mechanism once the databank is public.

### 5. E. Quality assurance and control of data

Quality assurance (QA) of all data is key to providing high quality and robust research findings (Ducharme et al., 2016). All QA steps of data collection (blinding of researchers, checking of data entry, standard operating procedures and calibration of equipment, particularly methods such as phantom scanning to describe scanner stability) should be recorded and provided with the data. Quality control should be implemented at all stages of the database from provenance to visualization. Data aggregation centres could provide a useful service with a common QC procedure across all datasets included. An important step is to be transparent on how this is implemented and what is tested on the data. If it is planned to incorporate processed data from external groups (e.g. templates, feature measures, etc.), how will the quality control be implemented? With increasing open access and secure web-based repositories, one option is to link to such repositories rather than incorporating secondary data into the database. This encourages early sharing of secondary (summary) data without the need to request access to the original databases (the model being pursued by UK Biobank) that have more stringent safeguards and comprehensive data access agreement to control research usage.

## 6. Database management

The legal and ethical framework of individual countries, and agreements reached between them, may affect how and where data are or can be stored. Systems are required to ensure data security, but allow appropriate access. Relevant approvals should be transparent, e.g. in publications and on websites.

During the meeting it was recognized that brain image databanks should have a Steering Committee, including independent and lay representatives, to monitor and review progress. This has the advantage of providing oversight of data usage and the opportunity to review data requests, but can be time consuming, it may be difficult to get agreement among stakeholders, and therefore delay access to data, and it can be difficult to identify lay representatives. Consideration should be given to how ‘sleuthing’ of individual’s data can be minimised or prevented, and to the legal and ethical aspects of data collection, storage and sharing. This is relevant both for adult and perinatal studies (for both mother and baby). Decisions need to be made in advance about what will happen to data if an individual decides to opt out; loses capacity in the future (e.g. develops dementia); or gains capacity to make his or her own decisions (e.g. a child growing up). Decisions should be able to be reviewed by the steering committee if legal or ethical frameworks change.

### 6. A. Legal and ethical issues

The legal framework may vary between countries, and this should be considered in international collaboration. The general principle is that researchers should satisfy the governing body, e.g. ethics committee, which they are processing and dealing with people’s data responsibly. It is useful to consider the concept of a ‘motivated intruder’: have reasonable steps been taken to protect data from someone making attempts to access the data.

Factors such as what will be done to divulge findings of abnormalities found on neuroimaging (e.g. tumours, or features of cerebrovascular disease, or multiple sclerosis) should be considered (Wardlaw et al., 2011; Wardlaw et al., 2015). Some primary studies have all scans evaluated by a neuroradiologist, others explicitly state that no feedback will be given on any tests. The issue for databanks is what to do if an abnormality of potential health significance is noted, or new health implications for existing findings come to light during secondary use – should the information be fed back to the participant, or the researchers that produced the data? These issues should be considered in the data donation agreement. If data are fully anonymised such feedback would not be possible.

The UK Health Research Authority (http://www.hra.nhs.uk/) is one example of a body that promotes research to improve clinical care of patients. The sharing and use of already collected data with appropriate safeguards fits this duty. The Royal Society, *Science as an Open Enterprise,* (2012) report, promotes *Intelligent Openness* – intelligibility, verifiability, accessibility if robust, if there is commercial confidence and personal privacy.

### 6. B. Publication

To maximise use, databanks should be registered on a publicly accessible registration website. Currently there is no international registry for neuroimaging studies or databanks. A general platform such as ClinicalTrials.gov could be used in the interim. One option would be a data posting website for imaging, like dbGaP for genetics

(http://www.ncbi.nlm.nih.gov/gap). Researchers should be encouraged to publish a ‘protocol paper’ which records how the databank was established, and the decisions made at each stage (e.g. for UK Biobank at http://www.ukbiobank.ac.uk/wp-content/uploads/2011/11/UK-Biobank-Protocol.pdf) and a ‘data paper’ that can include technical data and then be cited in the methods section of future results papers (e.g. for the imaging in UK Biobank http://www.nature.com/neuro/journal/v19/n11/full/nn.4393.html). The wording on citation and authorship should not contravene ICJME (International Committee of Medical Journal Editors) rules http://www.icmje.org/recommendations/browse/roles-and-responsibilities/defining-the-role-of-authors-and-contributors.html. There is also the possibility of having a Research Resource Identifier (RRID - Bandrowski et al., 2015). The consensus from our meeting was to discourage authorship, but to have the data paper cited along with the inclusion of the databank acronym with a web-reference or, better still, a DOI. A standardised template for reporting these papers (such as CONSORT for randomised controlled trials, see http://www.equator-network.org/) would be useful. This would allow an emphasis on clinical and epidemiological as well as technical perspectives.

### 6. C. Funding

Increasingly, funders are keen to encourage data sharing; indeed some go so far as to refuse future funding unless the results of prior funded studies have been published open source (http://www.wellcome.ac.uk/About-us/Policy/Policy-and-position-statements/WTD002766.htm). While most brain imaging studies are individually funded through grants to acquire data and undertake initial analyses, it has previously been difficult to obtain funding to create a brain image bank, or if initial funding was secured, then it may be insufficient to maintain the database long term, and deal with data requests. Storage and back up is, however, necessary, and inclusion for provision of this is encouraged. Researchers at ADNI estimate that 10-15% of funding, and 15% of time has been spent on data sharing (Wilhelm et al., 2014). The use of large-scale distributed computation can make the work more efficient, but users need to be aware of the heterogeneity of the constituent datasets e.g., in ENIGMA no one national government had to finance all the capital infrastructure. Similarly, a distributed image databank could be supported by funding from individual countries, much as are some multinational clinical trials. There is a tension between the desire to share the data and the feasibility of actually affording to do so, and a range of models exist. One model of cost-recovery used in UKBiobank (http://www.ukbiobank.ac.uk) is a fixed charge for reviewing the application, and a variable component depending on the data requested.

Some organisations, such as DataLad (http://datalad.org/) have developed infrastructure to provide access to scientific data available from various sources (e.g. lab or consortium websites or data sharing portals) through a single interface and integrated with software package managers

## Conclusions

Brain image biobanking is a rapidly evolving field. Several related and relevant projects will complement our recommendations, such as the International Neuroinformatics Coordinating Facility (INCF) Neuroimaging Data Sharing Task Force (wiki.incf.org/mediawiki/index.php/Neuroimaging_Task_Force) meeting held at Stanford University on January 27-30th 2015, which led to the development of the Brain Imaging Data Structure (BIDS - http://bids.neuroimaging.io/,Gorgolewski et al., 2016).

A federated international network of normative brain image biobanks is achievable (see table 2) and would have many advantages, including: 1) facilitating large scale meta-analyses of brain structure and function, in health and disease, following successful precedents (e.g. genetics – Hibar et al., 2015; depression - Schmaal et al., 2016; schizophrenia – van Erp et al., 2016; bipolar disorder – Hibar et al., 2016); 2) avoiding duplication of effort by data reuse, as occurs widely in physics; 3) providing population controls; 4) increasing research efficiency where research questions could be answered using existing data; and 5) providing a mechanism to replicate results in different demographic populations. The barriers to achieving this vision are political, ethical, technical, and financial, but a federated international group could work with funders, legal and ethical experts and industry along the lines of the solutions proposed here. Most large scale recent initiatives are still within geographical regions (e.g. the Obama BRAIN initiative; the European Human Brain Project), with some exceptions, such as the ENIGMA initiative spanning 35 countries (Thompson et al., 2015). Truly global, inter-regional initiatives are needed, to make full use of neuroimaging to understand the brain across the life-course.

---

**Table 2.** Action points for global data sharing via brain image databanks

| **Key issue** | **Reason** | **Suggested action** |
| --- | --- | --- |
| Data provenance unclear | Establish if results relevant for user's population | Clear data provenance tagged to each data item e.g. W3C-PROV |
| Acknowledge role of original studies, and databanks | Ensure clear recognition and ownership of data | Data use agreement templates should be published, and shared between databanks |
| Ensure appropriate informed consent | Define whether data storage, sharing and future use is permitted | Ethical committee review of applications, share consent templates between databanks |
| Oversight of databank | Appropriate use of data, data sharing | Studies should have a steering committee, with independent oversight, and consider legal/ethical/lay members |
| Variation in data collected | Difficult to share data, | Agree minimum dataset (e.g. |
|  | compare results | sex, age, handedness), and ontology (e.g. NIDM from INCF; BIDS) |
| Subject duplication | Avoid multiple entries from single individual | Global register of image banks; (or use brain image or genome as an identifier) |
| Databank changes with time | Unsure what data were in databank when analyses performed | Database version included with database; data accessed and study ID for included subjects provided for published papers |
| Volume of data | Need secure methods for data sharing | Safe Havens or Virtualization Desktop Infrastructure |
| Longevity of databank | Changes in hardware and software for imaging, data entry and storage | Involve technical experts in design, regular back up, including to remote location |
| Longevity of databank | Data storage, and sharing, as required by most funders of original projects | Funding required for database management and storage, and allowing open access |
| Large numbers of databanks | Can be difficult to find. Analysis of already-collected data may reduce research waste, compared to conducting new study | Register database e.g. on clinicaltrials.gov, or set up a register of brain imaging databases; early publication of protocol or data paper |
| Variability in databanks | Designed for wide range of purposes, may be used for other secondary aims | Standardised reporting guidelines for papers describing brain image databanks (such as CONSORT for randomised |
|  |  | controlled trials, see<br>http://www.equator-network.org/ ) |
| Authorship | Papers that use data from brain image databanks but where databank staff are not involved in writing the paper | Establish different authorship categories (as done for trials); reference to protocol or data paper |

## Appendix

Participants in meeting on ‘Development of human brain image banks and age-specific normative brain atlases’ held at Royal Society of Edinburgh, 28-29^th^ August 2014, which formed the basis for this paper:

Dr Trevor Ahearn, Dr Devasuda Anblagan, Dr John Ashburner, Ellen Backhouse, Marcelo Barria, Prof Richard Baldock, Dr Mark Bastin, Dr James Boardman, Prof Monique Breteler, Dr Albert Burger, Prof Jonathan Cavanagh, Dr James Cheney, James Cole, Prof Serena Counsell, Dr Simon Cox, Olimpia Curran, Dr David Alexander Dickie, Prof Klaus Ebmeier, Prof Alan Evans, Prof Nick Fox, Dr Lorna Gibson, Ludavica Griffanti, Dr Laura Gui, Lewis Hou, Dr Neda Jahanshad, Dr Dominic Job, Dr Rajeev Krishnadas, Dr Linda Lanyon, Prof Graeme Laurie, Prof Steve Lawrie, Prof Aad van der Lugt, Jennifer Macfarlane, Prof Andrew McIntosh, Dr Clare Mackay, Dr John McLean, Prof Paul Matthews, Prof Bernard Mazoyer, Prof Daniel Marcus, Shadia Mikhael, Prof Karla Miller, Prof Giovanni Montana, Prof Alison Murray, Dr Katie Nicol, Dr Thomas Nichols, Prof Wiro Niessen, Dr Mario Parra, Prof Mark Parsons, Dr Steve Pavis, Dr Cyril Pernet, Prof Jean-Baptiste Poline, Dr Douglas Potter, Prof Rebecca Reynolds, Prof Neil Roberts, Andrew Robson Prof Dave Robertson, Dr Mark Rodrigues, Dr David Rodriguez Gonzalez, Prof Burkhard Schafer, Prof Aziz Sheikh, Dr Ruth Sibbet, Dr Susan Shenkin, Dr Colin Smith, Prof Leslie Smith, Prof Steve Smith, Dr Roger Staff, Prof John Starr, Dr Amos Storkey, Daniel Taylor, Prof Paul Thompson, Dr Tony Stöcker, Dr Maria Valdes Hernandez, Dr Lana Vasung, Prof Joanna Wardlaw, Dr Heather Whalley, Dr Rebecca Woodfield, Prof David Wyper, Kaiming Yin, Dr May Yong, Dr Gabriel Ziegler.

## Acknowledgements

Thanks to Guarantors of Brain, British Geriatrics Society (Scottish Branch), Royal Society of Edinburgh, SINAPSE (Scottish Imaging Network: a Platform for Scientific Excellence) SPIRIT, International Neuroinformatics Coordinating Facility and Nuffield Foundation for contributions towards funding the meeting which formed the basis of this paper. TEN is supported by the Wellcome Trust (100309/Z/12/Z).

## Web references

http://www.erasmus-epidemiology.nl/research/ergo.htm Accessed 20^th^ July, 2016 https://www.dzne.de/en/research/research-areas/population-health-sciences/rhineland-study.html Accessed 20^th^ July, 2016

https://www.mcgill.ca/globalhealth/international-consortium-brain-mapping Accessed 20^th^ July, 2016

http://www.three-city-study.com/the-three-city-study.php Accessed 20^th^ July, 2016 http://www.imagen-europe.com/en/consortium.php Accessed 20^th^ July, 2016 http://www.wellcome.ac.uk/About-us/Policy/Policy-and-position-statements/WTD002766.htm Accessed 20^th^ July, 2016

http://www.icmje.org/recommendations/browse/roles-and-responsibilities/defining-the-role-of-authors-and-contributors.html Accessed 20^th^ July, 2016

http://mcin-cnim.ca/neuroimagingtechnologies/loris/ Accessed 20^th^ July, 2016

http://bids.neuroimaging.io/ Accessed 20^th^ July, 2016

http://bids.neuroimaging.io/http://nidm.nidash.org/ Accessed 20^th^ July, 2016

http://populationimaging.eu/ Accessed 20^th^ July, 2016

http://www.generationr.nl/ Accessed 20^th^ July, 2016

http://www.brainsimagebank.ac.uk/ Accessed 20^th^ July, 2016

http://www.dementiasplatform.uk/ Accessed 20^th^ July, 2016

http://www.oasis-brains.org/ Accessed 20^th^ July, 2016

http://www.humanconnectome.org/ Accessed 20^th^ July, 2016

http://www.gin.cnrs.fr/BILandGIN Accessed 20^th^ July, 2016

http://www.ukbiobank.ac.uk/ Accessed 20^th^ July, 2016

http://adni.loni.usc.edu/ Accessed 20^th^ July, 2016

http://enigma.ini.usc.edu/ Accessed 20^th^ July, 2016

http://www.humanbrainmapping.org/COBIDAS Accessed 20^th^ July, 2016

http://wp.doc.ic.ac.uk/dhcp Accessed 20^th^ July, 2016 http://cde.nih.gov/ Accessed 20^th^ July, 2016

http://www.w3.org/TR/prov-dm Accessed 20^th^ July, 2016 http://www.ncbi.nlm.nih.gov/gap Accessed 20^th^ July, 2016 http://www.equator-network.org/ Accessed 20^th^ July, 2016 http://www.hra.nhs.uk/ Accessed 20^th^ July, 2016 http://coins.mrn.org/ Accessed 20^th^ July, 2016 https://github.com/gbook/nidb Accessed 20^th^ July, 2016 http://scitran.github.io/ Accessed 20^th^ July, 2016 http://www.xnat.org/ Accessed 20^th^ July, 2016 http://dicom.nema.org/ Accessed 20^th^ July, 2016 http://www.w3.org/TR/prov-overview/ Accessed 20^th^ July, 2016 http://nidm.nidash.org Accessed 20^th^ July, 2016 https://www.humanbrainproject.eu/enGB Accessed 20^th^ July, 2016

http://www.neurovault.org Accessed 20^th^ July, 2016 https://www.openfmri.org Accessed 20^th^ July, 2016

